# AMBER: Assessment of Metagenome BinnERs

**DOI:** 10.1101/239582

**Authors:** Fernando Meyer, Peter Hofmann, Peter Belmann, Ruben Garrido-Oter, Adrian Fritz, Alexander Sczyrba, Alice C. McHardy

## Abstract

Reconstructing the genomes of microbial community members is key to the interpretation of shotgun metagenome samples. Genome binning programs deconvolute reads or assembled contigs of such samples into individual bins, but assessing their quality is difficult due to the lack of evaluation software and standardized metrics. We present AMBER, an evaluation package for the comparative assessment of genome reconstructions from metagenome benchmark data sets. It calculates the performance metrics and comparative visualizations used in the first benchmarking challenge of the Initiative for the Critical Assessment of Metagenome Interpretation (CAMI). As an application, we show the outputs of AMBER for ten different binnings on two CAMI benchmark data sets. AMBER is implemented in Python and available under the Apache 2.0 license on GitHub (https://github.com/CAMI-challenge/AMBER).

## Introduction

Metagenomics allows studying microbial communities and their members (populations) by shotgun sequencing. Evolutionary divergence and abundances of these members can vary widely, with genomes occasionally being very closely related to one another, representing strain-level diversity, or evolutionary far apart, whereas abundance can differ by several orders of magnitude. Genome binning software deconvolutes metagenomic reads or assembled sequences into bins representing genomes of the community members. A popular and performant approach in genome binning uses the covariation of read coverage and short k-mer composition of contigs with the same origin across co-assemblies of one or more related samples, though the presence of strain-level diversity substantially reduces bin quality [1].

Benchmarking methods for binning and other tasks in metagenomics, such as assembly and profiling, is crucial for both users and method developers. The former need to determine the most suitable programs and parametrizations for particular applications and data sets, and the latter need to compare their novel or improved method with existing ones. When lacking evaluation software or standardized metrics, both need to individually invest considerable effort in assessing methods. CAMI is a community-driven initiative aiming to tackle this problem by establishing evaluation standards and best practices, including the design of benchmark data sets and performance metrics [1,2]. Here, we describe AMBER, an evaluation package for the comparative assessment of genome binning reconstructions from metagenome benchmark data sets. It implements all metrics decided by the community to be most relevant for assessing the quality of genome reconstructions in the first CAMI challenge and is applicable to arbitrary benchmark data sets. AMBER automatically generates binning quality assessments outputs in flat files, as summary tables, rankings, and as visualizations in images and an interactive HTML page. It complements the popular CheckM software that assesses genome bin quality on real metagenome samples based on sets of single-copy marker genes [3].

## Methods

### Input

AMBER uses as input three types of files to assess binning quality for benchmark data sets: (1) a gold standard mapping of contigs or read IDs to underlying genomes of community members; (2) one or more files with predicted bin assignments for the sequences; and (3), a FASTA or FASTQ file with sequences. Benchmark metagenome sequence samples with a gold standard mapping can, for instance, be created with the CAMISIM metagenome simulator (https://github.com/CAMI-challenge/MetagenomeSimulationPipeline). A gold standard mapping can also be obtained for sequences (reads or contigs), provided that reference genomes are available, by aligning the sequences to these genomes. Popular read aligners are, for example, Bowtie [4] and BWA [5]. MetaQUAST [6] can also be used for contig alignment while it evaluates metagenome assemblies. High confidence alignments can then be used as mappings of the sequences to the genomes. The input files (1) and (2) use the Bioboxes binning format [7] (https://github.com/bioboxes/rfc/tree/master/data-format). AMBER also accepts as bin assignments individual FASTA files for each bin, as provided by MaxBin [8]. These can be converted to the Bioboxes format. Example files are provided in the AMBER GitHub repository (https://github.com/CAMI-challenge/AMBER).

### Metrics and accompanying visualizations

AMBER uses the gold standard mapping to calculate a range of relevant metrics [1] for one or more genome binnings of a given data set. We give below a more formal definition of all metrics than in [1], together with an explanation of their biological meaning.

#### Assessing the quality of bins

The purity and completeness, both ranging from 0 to 1, are commonly used measures for quantifying bin assignment quality, usually in combination [9]. We provide formal definitions below. As predicted genome bins have no label, e.g. a taxonomic one, the first step in calculating genome purity and completeness is **mapping each predicted genome bin to an underlying genome**. For this, AMBER uses one of the following choices:

(1) A predicted genome bin is mapped to the most abundant genome in that bin in number of base pairs. More precisely, let *X* be the set of predicted genome bins and *Y* the set of underlying genomes. We define a mapping of the predicted genome bin *x ∈ X* as *g*(*x*) *= y*, such that genome *y* maps to *x* and the overlap between *x* and *y*, in base pairs, is maximal among all *y ∈ Y*, i.e.

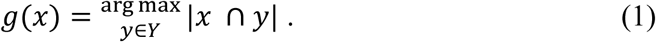
(2) A predicted genome bin is mapped to the genome whose largest fraction of base pairs has been assigned to the bin. In this case, we define a mapping *g′*(*x*) = *y* as

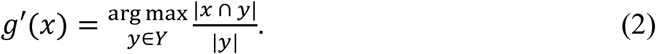

If more than a genome is completely included in the bin, i.e. | *x* ⋂ *y*|/|*y*| = 1.0 for more than a *y ∈ Y*, then the largest genome is mapped.

Using either option, each predicted genome bin is mapped to a single genome, but a genome can map to multiple bins or remain unmapped. Option 1 maps to each bin the genome that best represents the bin, since the majority of the base pairs in the bin belong to that genome, whereas option 2 maps to each bin the genome that best represents that genome, since most of the genome is contained in that specific bin. AMBER uses per default option 1. In the following, we use *g*\* to denote one of these mappings for simplicity whenever possible.

The **purity *p***, also known as precision, or specificity, quantifies the quality of genome bin predictions in terms of how trustworthy those assignments are. Specifically, the purity represents the ratio of base pairs originating from the mapped genome to all bin base pairs. For every predicted genome bin *x*,

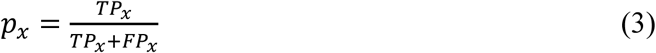

is determined, where the true positives *TP_x_* are the number of base pairs that overlap with the mapped genome *g*\*(*x*), i.e. *TP_x_* = |*x* ⋂ *g*\*(*x*)|, and the false positives *FP_x_* are the number of base pairs belonging to other genomes and incorrectly assigned to the bin. The sum *TP_x_* + *FP_x_* corresponds to the size of bin *x* in base pairs. See Figure 1 for an example of predicted genome bins and respective true and false positives.

**Figure 1:**
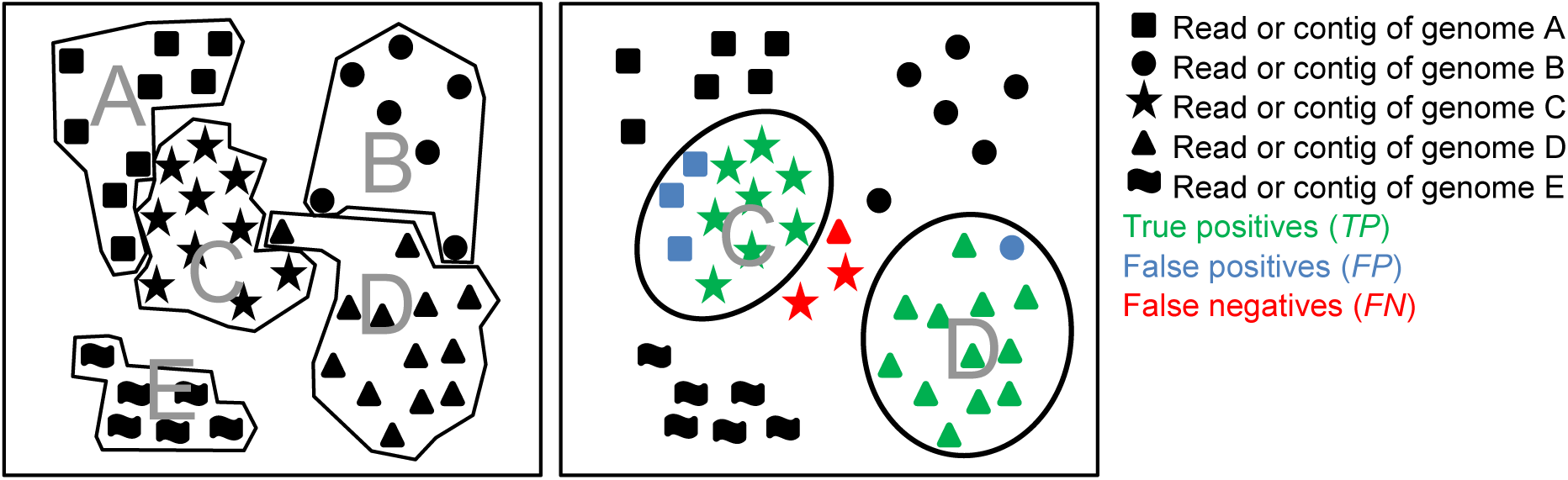
Schematic representation of establishing a bin-to-genome mapping for calculation of bin quality metrics. Reads and contigs of individual genomes are represented by different symbols and grouped by genome (left) or predicted genome bins (right). A bin-to-genome mapping is established using one of the criteria outlined in the text, with the upper bin mapping to genome C and the lower bin mapping to genome D. The mapping implies *TP*s*, FP*s and *FN*s for calculation of genome bin purity, completeness, contamination and overall sample assignment accuracy.

A related metric, the **contamination *c***, can be regarded as the opposite of purity and reflects the fraction of incorrect sequence data assigned to a bin (given a mapping to a certain genome). Usually, it suffices to consider either purity or contamination. It is defined for every predicted genome bin *x* as

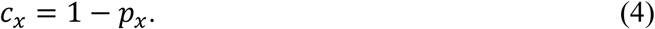

The **completeness *r***, also known as recall, or sensitivity reflects how complete a predicted genome bin is with regard to the sequences of the mapped underlying genome. For every predicted genome bin *x*,

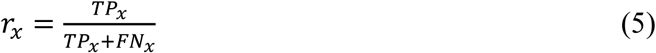

is calculated, where the false negatives *FN_x_* are the number of base pairs of the mapped genome *g*\*(*x*) that were classified to another bin or left unassigned. The sum *TP_x_* + *FN_x_* corresponds to the size of the mapped genome in base pairs.

Because multiple bins can map to the same genome, some bins might have a purity of 1.0 for a genome (if they exclusively contain its sequences), but the completeness for those bins sum up to at most 1.0 (if they include together all sequences of that genome). Genomes remaining unmapped are considered to have a completeness of zero and their purity is undefined.

As summary metrics, the **average purity** 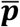 and **average completeness** 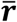 of all predicted genome bins can be calculated, which are also known in computer science as the macro-averaged precision and macro-averaged recall [10]. To these metrics, small bins contribute in the same way as large bins, differently from the sample-specific metrics discussed below. Specifically, the average purity 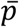 is the fraction of correctly assigned base pairs for all assignments to a given bin averaged over all predicted genome bins, where unmapped genomes are not considered. This value reflects how trustworthy the bin assignments are on average. Let *n_p_* = \*X*\ be the number of predicted genome bins. Then 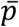 is calculated as

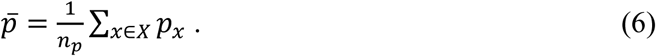

A related metric, the **average contamination** 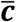 of a genome bin, is computed as

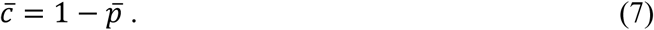

If very small bins are of little interest in quality evaluations, the **truncated average purity** 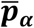 can be calculated, where the smallest predicted genome bins adding up to a specified percentage (the *α* percentile) of the data set are removed. For instance, the 99% truncated average purity can be calculated by sorting the bins according to their predicted size in base pairs and retaining all larger bins that fall into the 99% quantile, including (equally sized) bins that overlap the threshold. Let *S, S* ⊂ *X*, be the subset of predicted genome bins of *X* after applying the *α* percentile bin size threshold and *n_p_,_a_* = *\S\*. The truncated average purity 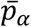 is calculated as

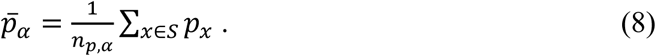

AMBER also allows to exclude other subsets of bins, such as bins representing viruses or circular elements.

While the average purity is calculated by averaging over all predicted genome bins, the average completeness 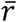 is averaged over all genomes, including those not mapped to genome bins (for which completeness is zero). More formally, let *Z* be the set of unmapped genomes, i.e. *Z* = {*y ∈ Y |* ∀*x ∈ X: g\**(*x*) *≠ y*}, and *n_r_* = |*X*| + |*Z*|, i.e. the sum of the number of predicted genome bins and the number of unmapped genomes. Then 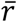 is calculated as

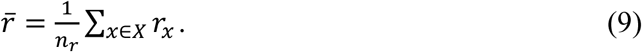

#### Assessing binnings of specific samples and in relation to bin sizes

Generally, it may not only be of interest how well a binning program does for individual bins, or all bins on average, irrespective of their sizes, but also how well it does overall for specific types of samples, where some genomes are more abundant than others. Binners may perform differently for abundant than for less abundant genomes, or for genomes of particular taxa, whose presences and abundances depend strongly on the sampled environment. To allow assessment of such questions, another set of related metrics exist, which either measure the binning performance for the entire sample, the binned portion of a sample, or to which bins contribute proportionally to their sizes.

To give large bins higher weight than small bins in performance determinations, the **average purity** 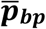 and **completeness** 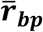 per base pair can be calculated as

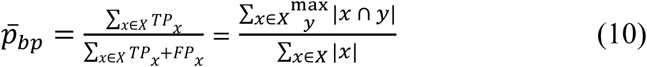

and

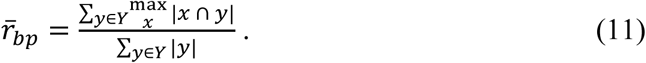

Equation (10) strictly uses the bin-to-genome mapping function *g*. Equation (11) computes the sum in base pairs of the intersection between each genome and the predicted genome bin that maximizes the intersection, averaged over all genomes. A genome that does not intersect with any bin results in an empty intersection. Binners achieving higher values of 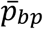 and 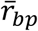 than for 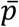 and 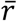 tend to do better for larger bins than for small ones, and for those with lower values it is the other way around.

The **accuracy *a*** measures the average assignment quality per base pair over the entire data set, including unassigned base pairs. It is calculated as

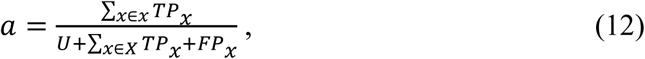

where *U* is the number of base pairs that were left unassigned. Like the average purity and completeness per base pair, large bins contribute more strongly to this metric than small bins.

Genome binners generate groups or clusters of reads and contigs for a given data set. Instead of calculating performance metrics established with a bin-to-genome mapping, another way to evaluate the quality of a clustering is to measure the similarity between the obtained and correct cluster partitions of the data set, corresponding here to the predicted genome bins and the gold standard contig or read genome assignments, respectively. This is accomplished with the Rand Index by comparing how pairs of items are clustered [11]. If two contigs or reads of the same genome are placed in the same predicted genome bin, these are here considered true positives *TP*. If two contigs or reads of different genomes are placed in different bins, these are considered true negatives *TN*. The Rand Index ranges from 0 to 1 and is the number of true pairs, *TP + TN*, divided by the total number of pairs. However, for a random clustering of the data set, the Rand Index would be larger than 0. The **Adjusted Rand Index** (ARI) corrects for this by subtracting the expected value for the Rand Index and normalizing the resulting value, such that the values still range from 0 to 1.

More formally, following [12], let *m* be the total number of base pairs assigned to any predicted genome bin and, *m_x,y_*, the number of base pairs of genome y assigned to predicted genome bin *x*. The ARI is computed as

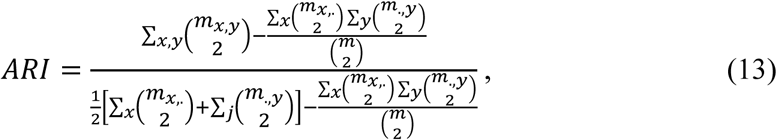

where 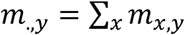 and 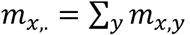. That is, 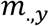 is the number of base pairs of genome *y* from all bin assignments and 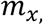 is the total number of base pairs in predicted genome bin *x*.

AMBER also provides ARI as a measure of assignment accuracy per sequence (contig or read) instead of per base pair by considering *m* to be the total number of sequences assigned to any bin and, *m_x,y_*, the number of sequences of genome *y* assigned to bin *x*. The meaning of 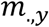 and 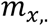 changes accordingly.

Importantly, the ARI is mainly designed for assessing a clustering of an entire data set, but some genome binning programs exclude sequences from bin assignment, thus assigning only a subset of the sequences from a given data set. If including this unassigned portion into the ARI calculation, the ARI becomes meaningless. AMBER, therefore, calculates the ARI only for the assigned portion of the data. For interpretation of these ARI values, the percentage of assigned data should also be considered (provided by AMBER together in plots).

## Output and visualization

AMBER combines the assessment of genome reconstructions from different binning programs or created with varying parameters for one program. The calculated metrics are provided as flat files, in several plots, and in an interactive HTML visualization. An example page is available at https://cami-challenge.github.io/AMBER/. The plots visualize:

- (Truncated) purity 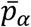 per predicted genome bin vs. average completeness 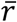 per genome, with the standard error of the mean
- Average purity per base pair 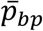 vs. average completeness per base pair 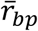
- Adjusted Rand Index ARI vs. percentage of assigned data
- Purity *p_x_* vs. completeness *r_x_* and boxplots for all predicted bins
- Heatmaps for individual binnings representing base pair assignments to predicted bins vs. their true origins from the underlying genomes

Heatmaps are generated from binnings without requiring a mapping, where rows represent the predicted genome bins and, columns, the genomes. The last row includes all unassigned base pairs for every individual genome and, individual entries, the number of base pairs assigned to a bin from a particular genome. Hence, the sum of all entries in a row corresponds to the bin size and, the sum of all column entries, to the size of the underlying genome. To facilitate the visualization of the overall binning quality, rows and columns are sorted as follows: for each predicted bin in each row, a bin-to-genome mapping function (*g*, per default) determines the genome (column) that maps to the bin and the true positive base pairs for the bin. Predicted bins are then sorted by the number of true positives in descending order from top to bottom in the matrix and genomes are sorted from left to right in the same order of the bin-to-genome mappings for the predicted bins. In this way, true positives concentrate in the main diagonal starting at the upper left corner of the matrix.

AMBER also provides a summary table with the number of genomes recovered with less than a certain threshold (5% and 10% per default) of contamination and more than another threshold (50%, 70%, and 90% per default) of completeness. This is one of the main quality measures used by CheckM [3] and in e.g. [13] and [14]. In addition, a ranking of different binnings by the highest average purity, average completeness, or the sum of these two metrics is provided as a flat file.

## Results

To demonstrate an application of AMBER, we performed an evaluation of the genome binning submissions to the first CAMI challenge, together with predictions from three more programs and new program versions, on two of the three challenge data sets. These are simulated benchmark data sets representing a single sample data set from a low complexity microbial community with 40 genomes and a 5-sample time series data set of a high complexity microbial community with 596 genome members. Both data sets include bacteria, the high complexity sample also archaea, high copy circular elements (plasmids and viruses) and substantial strain-level diversity. The samples were sequenced with paired-end 150 bp Illumina reads to a size of 15 GB for each sample. The assessed binners were CONCOCT [12], MaxBin 2.0.2 [8], MetaBAT [15], Metawatt 3.5 [16], and MyCC [17]. We generated results with newer program versions of MetaBAT and MaxBin. Furthermore, we ran Binsanity, Binsanity-wf [18], and COCACOLA [19] on the data sets. The commands and parameters used with the programs are available in the Supplementary information.

On the low complexity data set, MaxBin 2.2.4, as its previous version, performed very well, as did the new MetaBAT version (2.11.2, Figure 3, Supplementary Figure 1). On the high complexity data set, both MaxBin versions assigned less data than other programs, though with the highest purity (Figures 2, 3). MetaBAT 2.11.2 substantially improved over the previous version with all measures, recovering the most high quality bins and showing the highest interquartile range in the purity and completeness boxplots for the high complexity data set. MetaBAT 2.11.2 and MaxBin 2.0.2 also recovered the most genomes with more than specified thresholds of completeness and contamination on the high and the low complexity data sets, respectively (Table 1, Supplementary Table 1). Notably, some binners, such as CONCOCT, may require more than five samples for optimal performance. All results and evaluations are also available in the CAMI benchmarking portal (https://data.cami-challenge.org).

**Figure 2:**
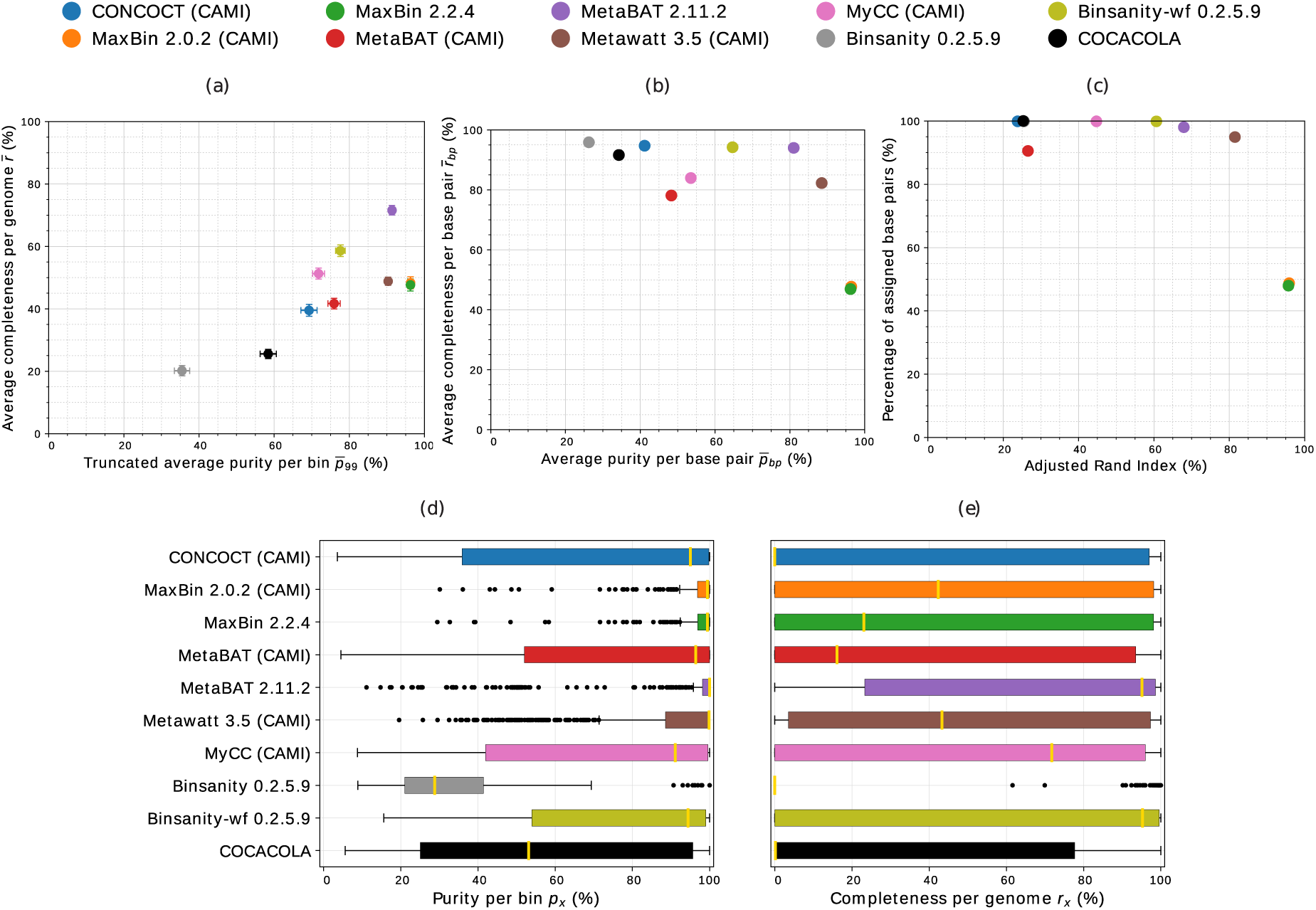
Assessment of genome bins reconstructed from CAMI’s high complexity challenge data set by different binners. Binner versions participating in CAMI are indicated in the legend in parentheses. (a) Average purity per bin (x-axis), average completeness per genome (y-axis), and respective standard errors (bars). As in the CAMI challenge, we report p_99_ with 1% of the smallest bins predicted by each program removed. (b) Average purity per base pair (x-axis) and average completeness per base pair (y-axis). (c) Adjusted Rand Index per base pair (x-axis) and percentage of assigned base pairs (y-axis). (d-e) Boxplots of purity per bin and completeness per genome, respectively.

**Figure 3:**
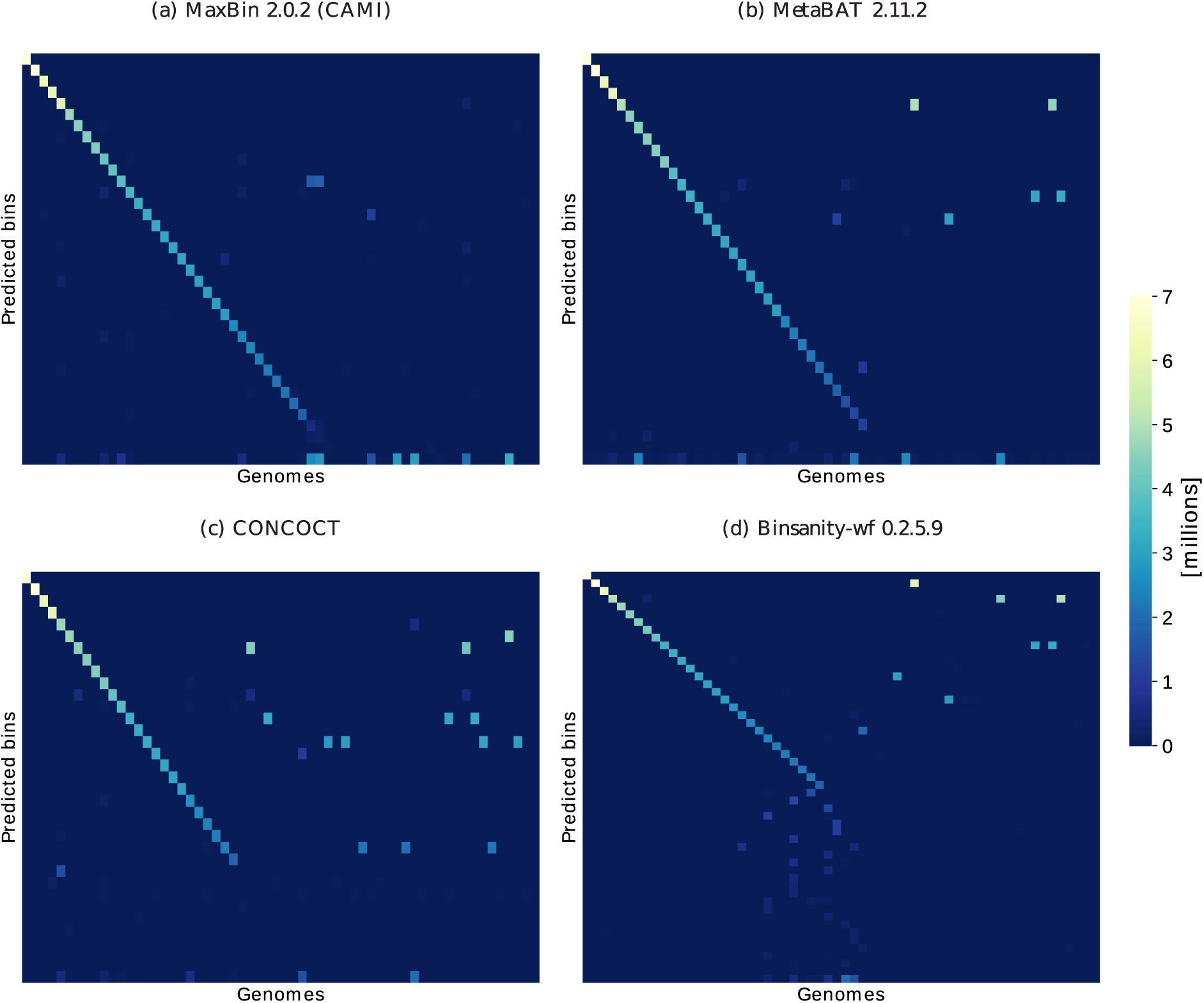
Heatmaps of confusion matrices for four different binnings for the low complexity data set of the first CAMI challenge representing the base pair assignments to predicted genome bins (y-axis) vs. their true origin from the underlying genomes or circular elements (x-axis). Rows and columns are sorted according to the number of true positives per predicted bin (see main text). Row scatter indicates a reduced average purity per base pair and thus underbinning (genomes assigned to one bin), whereas column scatter indicates a lower completeness per base pair and thus overbinning (many bins for one genome). The last row represents the unassigned bases per genome, allowing to assess the fraction of sample left unassigned. These views allow a more detailed inspection of binning quality relating to the provided quality metrics (Supplementary Figure 1).

**Table 1:** Respective number of genomes recovered from CAMI’s high complexity data set with less than 10% and 5% contamination and more than 50%, 70%, and 90% completeness.

| Genome binner<br>(% contamination) |  | Predicted bins<br>(% completeness) |  |  |
| --- | --- | --- | --- | --- |
|  |  | >50% | >70% | >90% |
| Gold standard |  | 596 | 596 | 596 |
| CONCOCT (CAMI) | <10% | 129 | 129 | 123 |
|  | <5% | 124 | 124 | 118 |
| MaxBin 2.0.2 (CAMI) | <10% | 277 | 274 | 244 |
|  | <5% | 254 | 252 | 224 |
| MaxBin 2.2.4 | <10% | 274 | 271 | 236 |
|  | <5% | 249 | 247 | 216 |
| MetaBAT (CAMI) | <10% | 173 | 152 | 126 |
|  | <5% | 159 | 140 | 118 |
| MetaBAT 2.11.2 | <10% | **427** | **417** | **361** |
|  | <5% | **414** | **404** | **353** |
| Metawatt 3.5 (CAMI) | <10% | 408 | 387 | 338 |
|  | <5% | 396 | 376 | 330 |
| MyCC (CAMI) | <10% | 189 | 182 | 145 |
|  | <5% | 166 | 159 | 127 |
| Binsanity 0.2.5.9 | <10% | 9 | 9 | 9 |
|  | <5% | 6 | 6 | 6 |
| Binsanity-refine 0.2.5.9 | <10% | 206 | 204 | 192 |
|  | <5% | 183 | 181 | 171 |
| COCACOLA | <10% | 88 | 87 | 75 |
|  | <5% | 69 | 69 | 60 |

## Conclusions

AMBER provides commonly used metrics for assessing the quality of metagenome binnings on benchmark data sets in several convenient output formats, allowing in-depth comparisons of binnings by different programs, software versions, or with varying parameter settings. As such, AMBER facilitates the assessment of genome binning programs on benchmark metagenome data sets, for bioinformaticians aiming to optimize data processing pipelines and method developers. The software is available as a standalone program, as a Docker image (automatically built with the provided Dockerfile), and in the CAMI benchmarking portal. We will continue to extend the metrics and visualizations according to community requirements and suggestions.

## Availability of supporting source code and requirements

Project name: AMBER: Assessment of Metagenome BinnERs

Project home page: https://github.com/CAMI-challenge/AMBER

Operating system(s): Platform independent

Programming language: Python 3.5

License: Apache 2.0

## Additional files

SupplementaryInformation.pdf

## Acknowledgements

The authors thank Christopher Quince for contributing Python code, all genome binning software developers participating in the CAMI challenge for their feedback on most relevant metrics, all developers who helped us to run their binning software, and the Isaac Newton Institute in Cambridge for its hospitality under the program MTG.

